# Human PGBD5 DNA transposase promotes site-specific oncogenic mutations in rhabdoid tumors

**DOI:** 10.1101/111138

**Authors:** Anton G. Henssen, Richard Koche, Jiali Zhuang, Eileen Jiang, Casie Reed, Amy Eisenberg, Eric Still, Ian C. MacArthur, Elias Rodríguez-Fos, Santiago Gonzalez, Montserrat Puiggròs, Andrew N. Blackford, Christopher E. Mason, Elisa de Stanchina, Mithat Gönen, Anne-Katrin Emde, Minita Shah, Kanika Arora, Catherine Reeves, Nicholas D. Socci, Elizabeth Perlman, Cristina R. Antonescu, Charles W. M. Roberts, Hanno Steen, Elizabeth Mullen, Stephen P. Jackson, David Torrents, Zhiping Weng, Scott A. Armstrong, Alex Kentsis

**Author notes:** Correspondence to: Alex Kentsis, MD, PhD. These authors contributed equally to this work (Anton G. Henssen, Richard Koche, Jiali Zhuang).

## Abstract

Genomic rearrangements are a hallmark of childhood solid tumors, but their mutational causes remain poorly understood. Here, we identify the *piggyBac transposable element derived 5* (*PGBD5*) gene as an enzymatically active human DNA transposase expressed in the majority of rhabdoid tumors, a lethal childhood cancer. Using assembly-based whole-genome DNA sequencing, we observed previously unknown somatic genomic rearrangements in primary human rhabdoid tumors. These rearrangements were characterized by deletions and inversions involving PGBD5-specific signal (PSS) sequences at their breakpoints, with some recurrently targeting tumor suppressor genes, leading to their inactivation. PGBD5 was found to be physically associated with human genomic PSS sequences that were also sufficient to mediate PGBD5-induced DNA rearrangements in rhabdoid tumor cells. We found that ectopic expression of *PGBD5* in primary immortalized human cells was sufficient to promote penetrant cell transformation *in vitro* and in immunodeficient mice *in vivo.* This activity required specific catalytic residues in the PGBD5 transposase domain, as well as end-joining DNA repair, and induced distinct structural rearrangements, involving PSS-associated breakpoints, similar to those found in primary human rhabdoid tumors. This defines PGBD5 as an oncogenic mutator and provides a plausible mechanism for site-specific DNA rearrangements in childhood and adult solid tumors.

## Introduction

Whole-genome analyses have now produced near-comprehensive topographies of coding mutations for certain human cancers, enabling both detailed molecular studies of cancer pathogenesis and potential of precisely targeted therapies ^1-5^. For certain childhood cancers, recent studies have begun to reveal the essential functions of complex non-coding structural variants that can induce aberrant expression of cellular proto-oncogenes ^6,7^. However, for many aggressive childhood cancers including embryonal tumors, such studies have identified distinct cancer subtypes that have no discernible coding mutations ^8-11^. In addition, while for some cancers, defects in DNA damage repair have been suggested to explain their increased incidence at a relatively young age, the causes of complex genomic rearrangements in cancers of young children without apparent widespread genomic instability remain largely unknown.

Rhabdoid tumor is a prototypical example of this question. Rhabdoid tumors occur in the developing tissues of infants and children, leading to tumors with neuroectodermal, epithelial and mesenchymal components in the brain, liver, kidney and other organs ^10,12,13^. Rhabdoid tumors that cannot be cured with surgery are generally chemotherapy resistant and almost uniformly lethal ^14^. Rhabdoid tumors exhibit inactivating mutations of *SMARCB1,* generally as a result of genomic rearrangements of the 22q11.2 chromosomal locus ^15^. These mutations can be inherited as part of the rhabdoid tumor predisposition syndrome, but are not thought to involve chromosomal instability ^13^. While *SMARCB1* mutations are sufficient to cause rhabdoid tumors in mice ^16^, human rhabdoid tumors have been observed to have multiple molecular subtypes and rearrangements of additional chromosomal loci that are poorly understood ^9,10,17,18^. These findings suggest that additional genetic elements and molecular mechanisms may contribute to the pathogenesis of rhabdoid tumors.

In humans, nearly half of the genome is comprised by sequences derived from transposons, including both autonomous and non-autonomous mobile genetic elements ^19^. The majority of human genes that encode enzymes that can mobilize transposons appear to be catalytically inactive, with the exception of L1 long interspersed repeated sequences (LINEs) that appear to induce structural genomic variation in human neurons and adenocarcinomas ^20-22^, *Mariner* transposase-derived SETMAR that functions in DNA repair ^23^, and *Transib-like* DNA transposase RAG1/2 that catalyzes somatic recombination of V(D)J receptor genes in lymphocytes ^24^. In particular, aberrant activity of RAG1/2 in lymphoblastic leukemias and lymphomas can induce the formation of chromosomal translocations that generate transforming fusion genes ^25-27^. The identity of and mechanisms by which similar genomic rearrangements may be formed in childhood solid tumors are unknown, but the existence of additional human recombinases that can induce somatic DNA rearrangements has long been hypothesized ^28^.

Recently, human PGBD5 and THAP9 have been found to catalyze transposition of synthetic DNA transposons in human cells ^29,30^. The physiologic functions of these activities are currently not known. PGBD5 is distinguished by its deep evolutionary conservation among vertebrates (~500 million years) and developmentally restricted expression in tissues from which childhood embryonal tumors, including rhabdoid tumors, are thought to originate ^30,31^. *PGBD5* is transcribed as a multi-intronic and non-chimeric transcript from a gene that encodes a full-length transposase that became immobilized on human chromosome 1 ^30,31^. Genomic transposition activity of PGBD5 requires distinct aspartic acid residues in its transposase domain, and specific DNA sequences containing inverted terminal repeats with similarity to the lepidopteran *Trichoplusia ni piggyBac* transposons ^30^. These findings, combined with the recent evidence that PGBD5 can induce genomic rearrangements that inactivate the *HPRT1* gene ^32^, prompted us to investigate whether PGBD5 may induce site-specific DNA rearrangements in human rhabdoid tumors that share developmental origin with cells that normally express *PGBD5.*

## Results

### Human rhabdoid tumors exhibit genomic rearrangements associated with PGBD5-specific signal sequence breakpoints

First, we analyzed the expression of PGBD5 in large, well-characterized cohorts of primary childhood and adult tumors (Supplementary Fig. 1a). We observed that *PGBD5* is highly expressed a variety of childhood and adult solid tumors, including rhabdoid tumors, but not in acute lymphoblastic or myeloid leukemias (Supplementary Fig. 1a). The expression of *PGBD5* in rhabdoid tumors was similar to that of embryonal tissues from which these tumors are thought to originate, and was not significantly associated with currently defined molecular subgroups or patient age at diagnosis (Supplementary Fig. 1a-f). To investigate potential PGBD5-induced genomic rearrangements in primary human rhabdoid tumors, we performed *de novo* structural variant analysis of whole-genome paired-end Illumina sequencing data for 31 individually-matched tumor versus normal paired blood specimens from children with extracranialrhabdoid tumors that are generally characterized by inactivating mutations of *SMARCB1* ^10^. By virtue of their repetitive nature, sequences derived from transposons present challenges to genome analysis. Thus, we reasoned that genome analysis approaches that do not rely on short-read alignment algorithms, such as the local assembly-based algorithm laSV and the tree-based sequence comparison algorithm SMuFin might reveal genomic rearrangements that otherwise might escape conventional algorithms ^33,34^.

Using this assembly-based approach, we observed recurrent rearrangements of the *SMARCB1* gene on chromosome 22q11 in nearly all cases examined, consistent with the established pathogenic function of inactivating mutations of *SMARCB1* in rhabdoid tumorigenesis (Fig. 1a). In addition, we observed previously unrecognized somatic deletions, inversions and translocations involving focal regions of chromosomes 1, 4, 5, 10, and 15 (median = 3 per tumor), which were recurrently altered in more than 20% of cases (Fig. 1a, Data S1). These results indicate that in addition to the pathognomonic mutations of *SMARCB1,* human rhabdoid tumors are characterized by additional distinct and recurrent genomic rearrangements.

**Fig 1.**
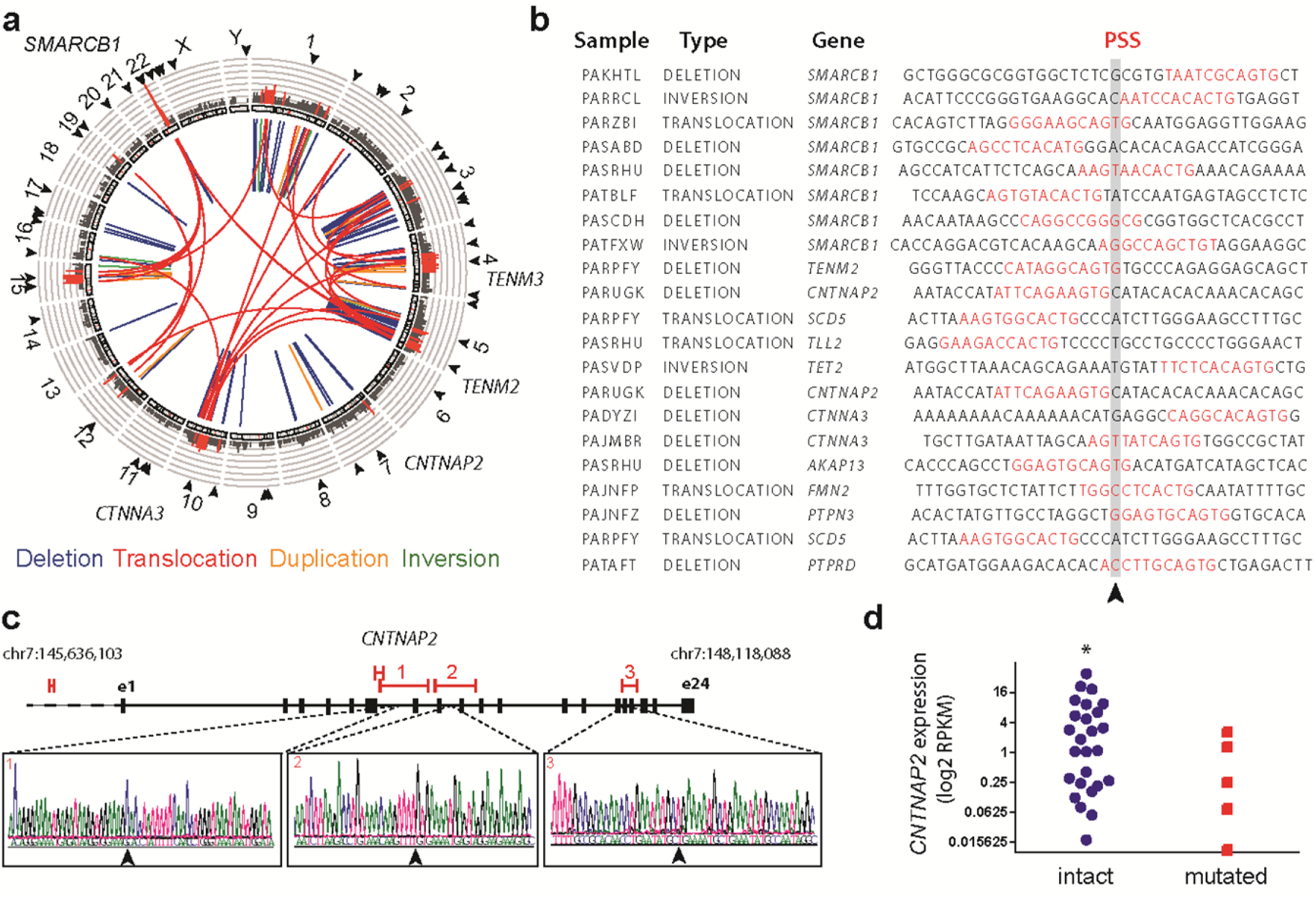
Human rhabdoid tumors exhibit genomic rearrangements associated with PGBD5-specific signal sequence breakpoints. **(a)** Aggregate Circos plot of somatic structural variants identified in 31 human rhabdoid tumors using laSV, as marked for PSS-containing breakpoints (outer ring, arrowheads), recurrence (middle ring histogram, rearrangements occurring in >3 out of 31 samples and highlighted in red for rearrangements with recurrence frequency greater than 13%), and structural variant type (inner lines, as color-labeled). Recurrently rearranged genes are labeled. **(b)** Representation of 21 structural variant breakpoints in rhabdoid tumors identified to harbor PSS sequences (red) within 10 bp of the breakpoint junction (arrowhead). **(c)** Recurrent structural variants of *CNTNAP2* (red) with gene structure (black) and Sanger sequencing of the rearrangement breakpoints. **(d)** *CNTNAP2* mRNA expression in primary rhabdoid tumors as measured using RNA sequencing in *CNTNAP2* mutant (red) as compared to *CNTNAP2* intact (blue) specimens (\**p =* 0.017 by t-test for intact vs. mutant *CNTNAP2).*

To determine whether any of the observed genomic rearrangements may be related to PGBD5 DNA transposase or recombinase activity, we first used a forward genetic screen to identify PGBD5-specific signal (PSS) sequences that were specifically found at the breakpoints of PGBD5-induced deletions, inversions and translocations that caused inactivation of the *HPRT1* gene in a thioguanine resistance assay ^32^. Using these PSS sequences as templates for supervised analysis of the somatic genomic rearrangements in primary human rhabdoid tumors, we identified specific PSS sequences associated with the breakpoints of genomic rearrangements in rhabdoid tumors (*p* = 1.1 × 10^−10^, hypergeometric test; Fig. 1b, Supplementary Fig. 2). By contrast, we observed no enrichment of the RAG1/2 recombination signal (RSS) sequences at the breakpoints of somatic rhabdoid tumor genomic rearrangements, in spite of their equal size to PSS sequences, consistent with the lack of *RAG1/2* expression in rhabdoid tumors. Likewise, we did not find significant enrichment of PSS motifs at the breakpoints of structural variants and genomic rearrangements in breast carcinomas that lack *PGBD5* expression, even though these breast carcinoma genomes were characterized by high rates of genomic instability (Data S1). In total, 580 (52%) out of 1121 somatic genomic rearrangements detected in rhabdoid tumors contained PSS sequences near the rearrangement breakpoints (Data S1).

Overall, the majority of the observed rearrangements were deletions and translocations (Fig. 1a, Supplementary Fig. 3a). Notably, we found recurrent PSS-containing genomic rearrangements affecting the *CNTNAP2, TENM2, TENM3,* and *TET2* genes (Fig. 1a-c, Supplementary Fig. 3c, Data S1). Using allele-specific polymerase chain reaction (PCR) followed by Sanger DNA sequencing, we confirmed three of the observed PGBD5-induced intragenic *CNTNAP2* deletions and rearrangement breakpoints (Fig. 1c). Likewise, we confirmed somatic nature of mutations of *CNTNAP2* and *TENM3* by allele-specific PCR in matched tumor and normal primary patient specimens (Supplementary Fig. 3d-h).

*CNTNAP2*, a member of the neurexin family of signaling and adhesion molecules, has been previously found to function as a tumor suppressor gene in gliomas ^35^. Consistent with the potential pathogenic functions of the apparent PGBD5-induced *CNTNAP2* rearrangements in rhabdoid tumors found in our analysis, *CNTNAP2* has also been recently reported to be recurrently deleted in an independent cohort of rhabdoid tumor patients ^18^. By using comparative RNA sequencing gene expression analysis, we found that recurrent genomic rearrangements of *CNTNAP2* in our cohort were indeed associated with significant reduction of its mRNA transcript expression in genomically rearranged primary cases as compared to those lacking *CNTNAP2* rearrangements (*p* = 0.017, t-test; Fig. 1d). Additional mechanisms, including as of yet undetected mutations or silencing ^35^, may contribute to the loss of *CNTNAP2* expression in apparently non-rearranged cases (Fig. 1d).

Interestingly, some of the observed genomic rearrangements with PSS-containing breakpoints in rhabdoid tumors involved *SMARCB1* deletions (Fig. 1a-b, Data S1), suggesting that in a subset of rhabdoid tumors, PGBD5 activity may contribute to the somatic inactivation of *SMARCB1* in rhabdoid tumorigenesis. Similarly, we observed recurrent interchromosomal translocations and complex structural variants containing breakpoints with the PSS motifs that involved *SMARCB1* (Fig. 1b, Data S1), including chromosomal translocations, previously observed using cytogenetic methods ^17^. For example, we verified the t(5;22) translocation using allele-specific PCR followed by Sanger sequencing of the translocation breakpoint (Suppl. Fig. 3i-j). In all, these results indicate that human rhabdoid tumors exhibit recurrent complex genomic rearrangements that are defined by PSS breakpoint sequences specifically associated with PGBD5, at least some of which appear to be pathogenic and may be coupled with inactivating mutations of *SMARCB1* itself.

### PGBD5 is physically associated with human genomic PSS sequences that are sufficient to mediate DNA rearrangements in rhabdoid tumor cells

In prior studies, human PGBD5 has been found to localize to the cell nucleus ^31^. To test whether PGBD5 in rhabdoid tumor cells is physically associated with genomic PSS-containing sequences, as would be predicted for a DNA transposase that induces genomic rearrangements, we used chromatin immunoprecipitation followed by DNA sequencing (ChIP-seq) to determine the genomic localization of endogenous PGBD5 in human G401 rhabdoid tumor cells. We observed that human DNA regions bound by PGBD5 were significantly enriched for PSS motifs (*p* = 2.9 × 10^−29^,hypergeometric test), in contrast to the scrambled PSS sequences of identical composition, or the functionally unrelated RSS sequences of equal size that showed no significant enrichment (*p* = 0.28 and 1.0, respectively, hypergeometric test; Fig. 2a).

**Fig. 2.**
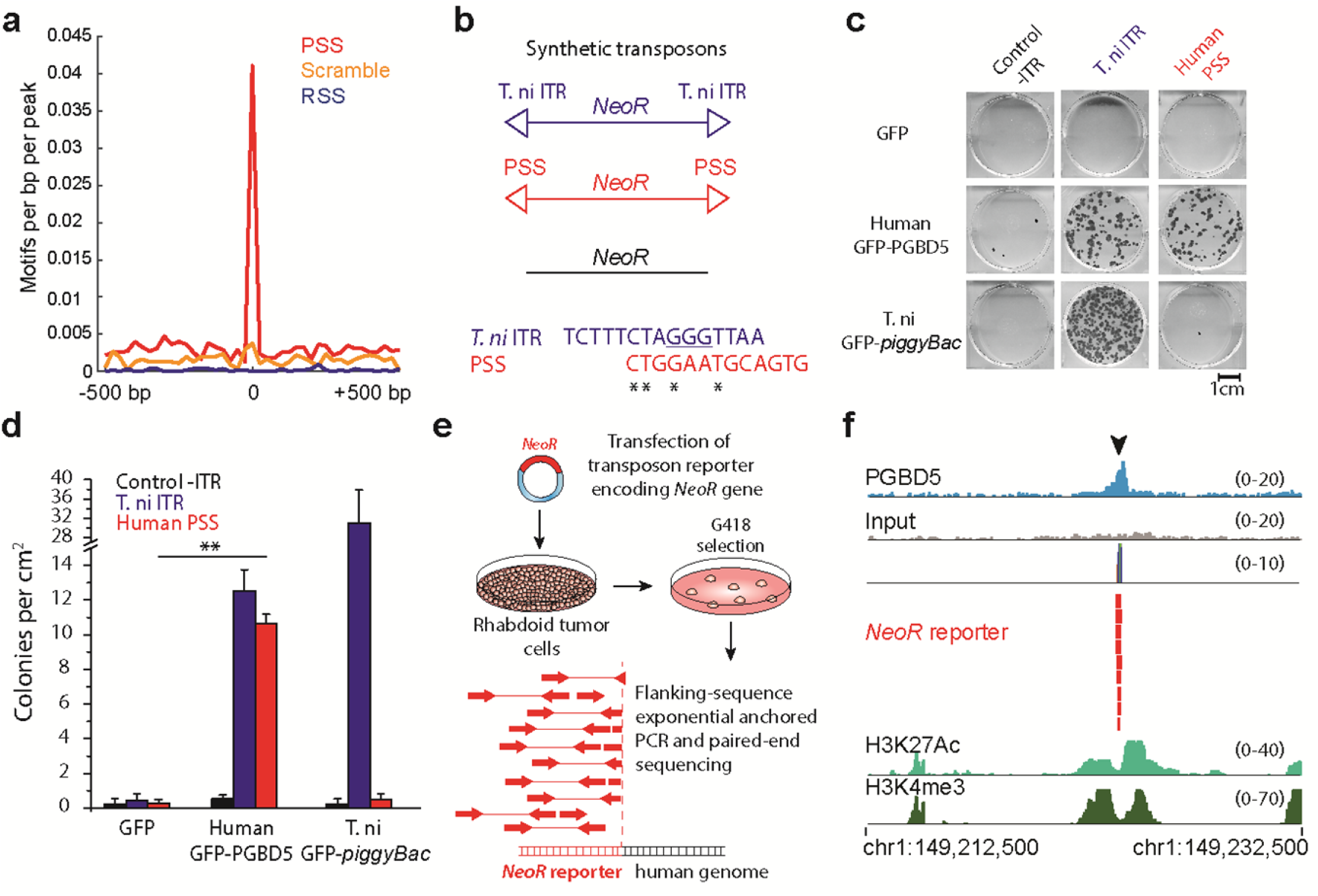
PGBD5 is physically associated with human genomic PSS sequences that are sufficient to mediate DNA rearrangements in rhabdoid tumor cells. **(a)** Genomic distribution of PGBD5 protein in G401 rhabdoid tumor cells as a function of enrichment of PSS (red) as compared to scrambled PSS (orange) and RAG1 recombination signal sequence (RSS, blue) controls as measured using PGBD5 ChIP-seq(*p*= 2.9 x 10^-29^ for PSS, *p*= 0.28 for scrambled PSS, *p*= 1.0 for RSS by hypergeometric test). **(b)** Schematic of synthetic transposon substrates used for DNA transposition assays, including transposons with *T. ni* ITR marked by triangles in blue, transposons with PGBD5-specific signal sequence (PSS) marked by triangles in red and transposons lacking ITRs marked in black (top) and sequence alignment of *T. ni* ITR compared to human PSS (bottom). **(c)** Representative photographs of Crystal violet-stained colonies obtained upon G418 selection in the transposon reporter assay. **(d)** Genomic DNA transposition assay as measured using neomycin resistance clonogenic assays in HEK293 cells co-transfected with human *GFP-PGBD5* or control *GFP* and *T.ni GFP-PiggyBac,* and transposon reporters encoding the neomycin resistance gene flanked by human PSS (red), as compared to control reporters lacking inverted terminal repeats (-ITR, black) and *T. nipiggyBac* ITR (blue). ** *p* = 5.0 x 10^-5^.Lepidopteran *T. ni* PiggyBac DNA transposase and its *piggyBac* ITR serve as specificity controls. Errors bars represent standard deviations of three biological replicates. **(e)**Schematic model of transposition reporter assay in G401 rhabdoid tumor cells followed by flanking sequence exponential anchored-polymerase chain reaction (FLEA-PCR) and Illumina paired-end sequencing. **(f)** Genomic integration of synthetic *NeoR* transposons (red) by endogenous PGBD5 in G401 rhabdoid tumor cells at PSS site (arrowhead), as shown in the ChIP-seq genome track of PGBD5 (blue), as compared to its sequencing input (gray), and H3K27Ac and H3K4me3 (bottom), consistent with the bound PGBD5 transposase protein complex.

To test the hypothesis that PGBD5 can act directly on human PSS-containing DNA sequences to mediate their genomic rearrangements, we used the previously established DNA transposition reporter assay ^30^. Human embryonic kidney (HEK) 293 cells were transiently transfected with plasmids expressing human *GFP-PGBD5,* hyperactive lepidopteran *T. ni GFP-PiggyBac* DNA transposase or control *GFP*, in the presence of reporter plasmids encoding the neomycin resistance gene (*NeoR*) flanked by a human PSS sequence, as identified from rhabdoid tumor rearrangement breakpoints (Suppl. Fig. 2-3, Data S1), lepidopteran *piggyBac* inverted terminal repeat (ITR) transposon sequence ^30^, or control plasmids lacking flanking transposon elements (Fig. 2b). Clonogenic assays of transfected cells in the presence of G418 to select neomycin resistant cells with genomic reporter integration demonstrated that GFP-PGBD5, but not control GFP, exhibited efficient activity on reporters containing terminal repeats with the human PSS sequences, but not control reporters lacking flanking transposon elements (*p* = 5.0 x 10^-5^, t-test; Fig. 2c & d). This activity was specific since the lepidopteran GFP-PiggyBac DNA transposase, which can efficiently mobilize its own *piggyBac* transposons, did not mobilize reporter plasmids containing human PSS sequences (Fig. 2c & d).

To determine whether endogenous PGBD5 can mediate genomic rearrangements in rhabdoid cells, we transiently transfected human G401 rhabdoid cells with the neomycin resistance gene transposon reporter plasmids, and determined their chromosomal integrations by using flanking sequence exponential anchored (FLEA) PCR to amplify and sequence specific segments of the human genome flanking transposon integration sites (Fig. 2e, Supplementary Fig. 4) ^30^. Similar assays in HEK293 cells that lack *PGBD5* expression fail to induce measureable genomic integration of reporter transposons (Fig. 2c & d). In contrast, we observed that endogenous PGBD5 in G401 rhabdoid tumor cells was sufficient to mediate integrations of transposon-containing DNA into human genomic PSS-containing sites (Fig. 2f, Supplementary Tables 5 & 6). This activity was specifically observed for transposon reporters with intact transposons, but not those in which the essential 5’-<u>GGG</u>TTAACCC-3’ hairpin structure was mutated to 5’-<u>ATA</u>TTAACCC-3’ (Supplementary Table 5) ^30^. Thus, PGBD5 physically associates with human genomic PSS sequences that are sufficient to mediate DNA rearrangements of synthetic reporters in rhabdoid tumor cells.

### PGBD5 expression in genomically stable primary human cells is sufficient to induce malignant transformation *in vitro* and *in vivo*

Recurrent somatic genomic rearrangements in primary rhabdoid tumors associated with PGBD5-specific signal sequence breakpoints, their targeting of tumor suppressor genes, and specific activity as genomic rearrangement substrates raise the possibility that PGBD5 DNA transposase activity may be sufficient to induce tumorigenic mutations that contribute to malignant cell transformation. To determine if PGBD5 can act as a human cell transforming factor, we used established transformation assays of primary human foreskin BJ and retinal pigment epithelial (RPE) cells immortalized with telomerase ^36^. Primary RPE and BJ cells at passage 3-5 can be immortalized by the expression of human *TERT* telomerase *in vitro,* undergo growth arrest upon contact inhibition, and fail to form tumors upon transplantation in immunodeficient mice *in vivo* ^36^. Prior studies have established the essential requirements for their malignant transformation by the concomitant dysregulation of P53, RB, and RAS pathways ^36^. Thus, transformation of primary human RPE and BJ cells enables detailed studies of human PGBD5 genetic mechanisms that cannot be performed using mouse or other heterologous model systems.

To test whether *PGBD5* has transforming activity in human cells, we used lentiviral transduction to express *GFP-PGBD5* and control *GFP* transgenes in telomerase-immortalized RPE and BJ cells, at levels that are 1.1-5 and 1.5-8 fold higher as compared to primary rhabdoid tumor specimens and cell lines, respectively (Fig. 3a & b). We observed that *GFP-PGBD5*-expressing but not non-transduced or *GFP*-expressing RPE and BJ cells formed retractile colonies in monolayer cultures and exhibited anchorage-independent growth in semisolid cultures, a hallmark of cell transformation (Fig. 3c & d). When transplanted into immunodeficient mice, *GFP-PGBD5*-expressing RPE and BJ cells formed subcutaneous tumors with similar latency and penetrance to that seen in cells expressing both mutant *HRAS* and the SV40 large T antigen that dysregulates both P53 and RB pathways (LTA; Fig. 3f & g, Supplementary Figure 5). Importantly, both RPE and BJ cells transformed by GFP-PGBD5 had stable, diploid karyotypes when passaged *in vitro* (Supplementary Figure 6). By contrast, expression of the distantly related lepidopteran *GFP-PiggyBac* DNA transposase which exerts specific and efficient transposition activity on lepidopteran *piggyBac* transposon sequences (Fig. 2d), failed to transform human RPE cells (Fig. 3e), in spite of being equally expressed (Supplementary Fig. 7a). These results indicate that the PGBD5 transposase can specifically transform human cells in the absence of chromosomal instability both *in vitro* and *in vivo.*

**Fig.3.**
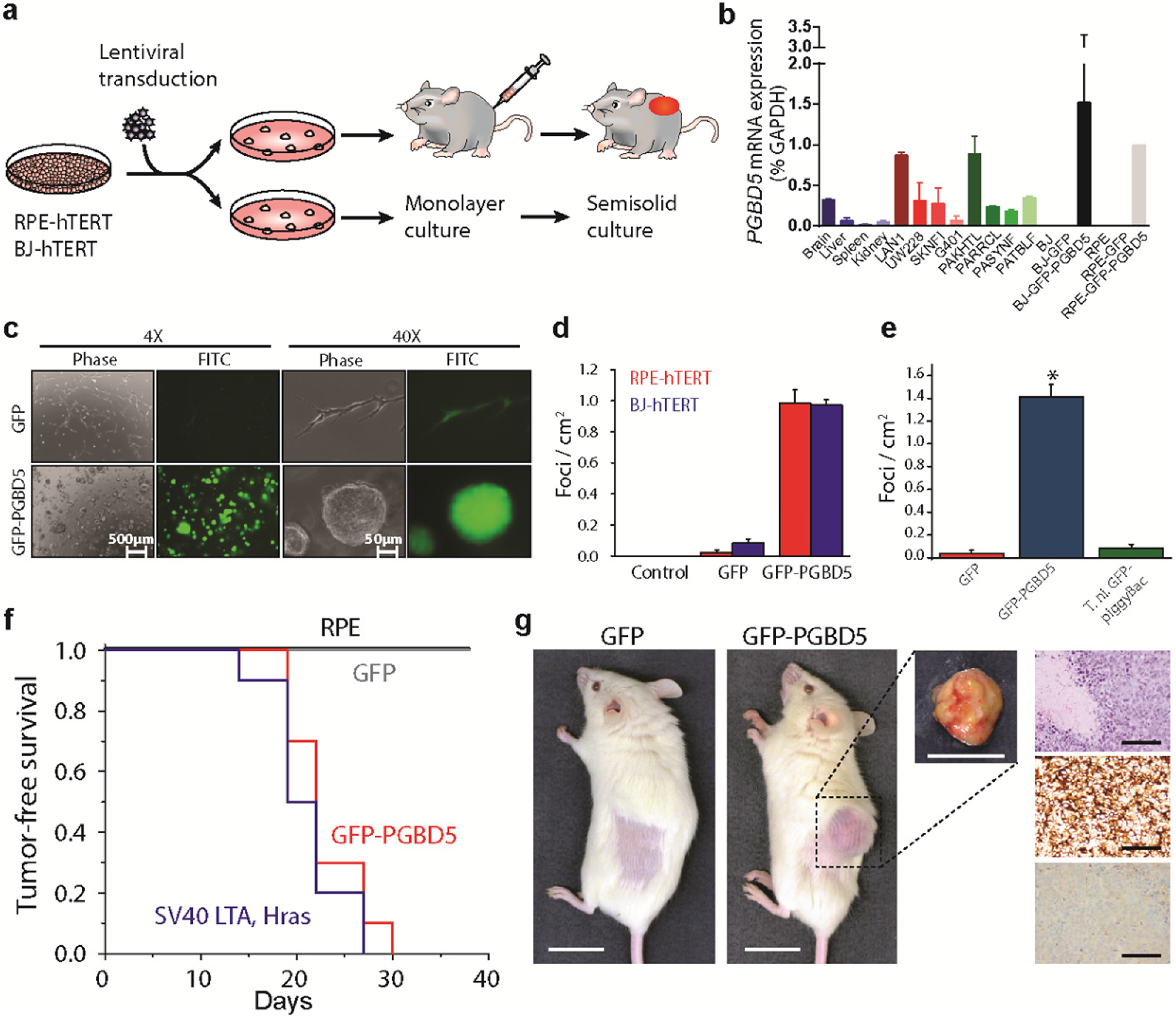
Ectopic expression of *PGBD5* in human cells leads to oncogenic transformation both *in vitro* and *in vivo.* **(a)**Schematic for testing transforming activity of PGBD5. **(b)** Relative *PGBD5* mRNA expression measured by quantitative RT-PCR in normal mouse tissues (brain, liver, spleen and kidney), as compared to human tumor cell lines (rhabdoid G401, neuroblastoma LAN1 and SK-N-FI, medulloblastomaUW-228 cells), primary human rhabdoid tumors (PAKHTL, PARRCL, PASYNF, PATBLF), and BJ and RPE cells stably transduced with *GFP-PGBD5* and *GFP.* Error bars represent standard deviations of 3 biological replicates. **(c)** Representative images of GFP-PGBD5-transduced RPE cells grown in semisolid media after 10 days of culture, as compared to control GFP-transduced cells. **(d)** Number of refractile foci formed in monolayer cultures of RPE and BJ cells expressing *GFP-PGBD5* or *GFP,* as compared to non-transduced cells (*p*= 3.6 x 10^-5^ and 3.9 x 10^-4^ for *GFP-PGBD5* vs. *GFP* for BJ and RPE cells, respectively). **(e)** Expression of *T. ni GFP-PiggyBac* does not lead to the formation of anchorage independent foci in monolayer culture (\**p*= 3.49 x 10^-5^ for *GFP-PGBD5* vs. *T. ni GFP-PiggyBac).* Error bars represent standard deviations of 3 biological replicates. **(f)** Kaplan-Meier analysis of tumor-free survival of mice with subcutaneous xenografts of RPE cells expressing *GFP-PGBD5* or *GFP* control, as compared to non-transduced cells or cells expressing SV40 large T antigen (LTA) and *HRAS (n*= 10 mice per group, *p*< 0.0001 by log-rank test). **(g)** Representative photographs (from left) of mice with shaved flank harboring RPE xenografts (scale bar = 1 cm). Tumor excised from mouse harboring *GFP-PGBD5* expressing tumor (scale bar = 1 cm). Photomicrograph of *GFP-PGBD5* expressing tumor (top to bottom: hematoxylin and eosin stain, vimentin, and cytokeratin, scale bar = 1 mm).

### PGBD5-induced cell transformation requires 274 DNA transposase activity

To test whether the cell transforming activity of PGBD5 requires its transposase enzymatic activity, we used PGBD5 point mutants that are proficient or deficient in DNA transposition in reporter assays ^30^. Thus, we compared E373A and E365A PGBD5 mutants that retain wild-type transposition activity ^30^, to D168A, D194A, D386A or their double D194A/D386A (DM) and triple D168A/D194A/D386A (TM) mutants that occur on residues required for efficient DNA transposition *in vitro,* consistent with their evolutionary conservation and putative function as the DDD/E catalytic triad for the phosphodiester bond hydrolysis reaction ^30^. After confirming stable and equal expression of these PGBD5 mutants in RPE cells by Western immunoblotting (Fig. 4a), we assessed their transforming activity with contact inhibition assays in monolayer cultures and transplantation in immunodeficient mice. Whereas ectopic expression of wild-type GFP-PGBD5 induced efficient and fully penetrant cell transformation, neither D168A, nor D194A, nor DM or TM mutants deficient in transposition function in reporter assays induced contact inhibition *in vitro* or tumor formation *in vivo* (Fig. 4b & d). By contrast, transposition-proficient E373A and E365A mutants exhibited the same transforming activity as wild-type GFP-PGBD5 (Fig. 4b and 4d). Importantly, we confirmed that the catalytic mutants of GFP-PGBD5 on average retained their chromatin localization as compared to wild-type PGBD5, as assessed using ChIP-seq (Fig. 4c). Although the D386A mutant exhibited reduced transposition activity in reporter assays *in vitro*^30>^, its expression induced wild-type transforming activity *in vivo* (Fig. 4d). This suggests that the transforming activity of PGBD5 may involve non-canonical DNA transposition or recombination reactions, consistent with the dispensability of some catalytic residues for certain type of DNA transposase-induced DNA rearrangements ^37,38^. Thus, cell transformation induced by PGBD5 requires its nuclease activity.

**Fig. 4.**
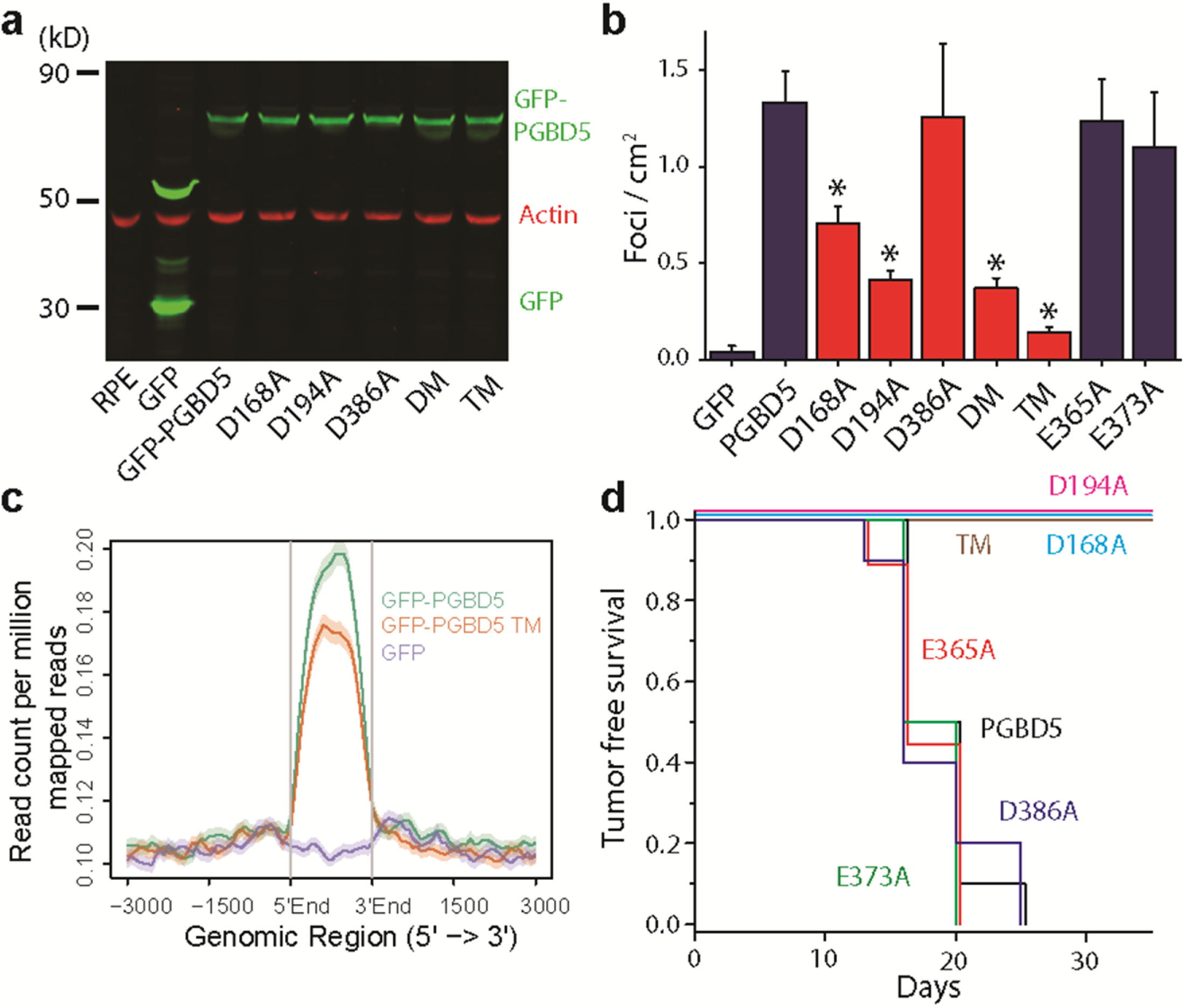
PGBD5 transposase activity is necessary to transform human cells. **(a)** Western blot of GFP in RPE cells expressing *GFP-PGBD5, GFP-PGBD5* mutants, and *GFP* compared to RPE cells (DM = double mutant D194A/D386A; TM = triple mutant D168A/D194A/D386A). **(b)** Number of refractile foci formed in monolayer culture in RPE and BJ cells stably expressing *GFP-PGBD5* or control *GFP*, as compared to non-transduced cells and cells expressing *GFP-PGBD5* mutants (red =transposase deficient mutants, blue =transposase proficient mutants, * *p* = 2.1 × 10^−4^ for *D168A* vs. *GFP-PGBD5, p* = 2.7 × 10^−6^ for *D194A* vs. *GFP-PGBD5, p* = 1.8 × 10^−6^ for *D194A/D386A* vs. *GFP-PGBD5, p* = 2.4 × 10^−7^ for *D168A/D194A/D386A* vs. *GFP-PGBD5).* Error bars represent standard deviations of three biological replicates. **(c)** Composite plot of ChIP-seq of GFP-PGBD5 (green), as compared to the GFP-PGBD5 D168A/D194A/D386A catalytic TM mutant (orange) and GFP control (purple). **(d)** Kaplan-Meier analysis of tumor-free survival of mice with subcutaneous xenografts of RPE cells expressing *GFP-PGBD5* as compared to cells expressing *GFP-PGBD5* mutants (n = 10 per group, *p*< 0.0001 by log-rank test).

### Transient expression of PGBD5 is sufficient for PGBD5-induced cell transformation

If PGBD5 can induce transforming genomic rearrangements, then transient exposure to PGBD5 should be sufficient to heritably transform human cells. To test this prediction, we generated doxycycline-inducible *PGBD5*-expressing RPE cells, and using Western immunoblotting confirmed lack of detectable expression of the enzyme in the absence of doxycycline and its induction upon exposure to doxycycline *in vitro* (Supplementary Fig. 7b). When transplanted into immunodeficient mice whose doxycycline chow treatment (–Dox) was stopped upon macroscopic signs of tumor formation (Fig. 5a, Supplementary Fig. 7c), the transduced cells retained essentially the same tumorigenicity as seen in continuously treated (+Dox) animals or in those transplanted with constitutively expressing *GFP-PGBD5* cells (Supplementary Fig. 7c). Importantly, we confirmed the absence of measureable PGBD5 expression in tumors harvested from –Dox animals by Western immunoblotting (Fig. 5a, inset). Consistent with cell transformation by transient expression of *PGBD5,* both–Dox and +Dox tumors were indistinguishable histopathologically (Fig. 5b). To investigate the potential irreversibility and heritability of cell transformation induced by transient PGBD5 expression, we transplanted tumors harvested from–Dox and+Dox animals into secondary recipients, and observed that tumors were induced with the same latency and penetrance in both –Dox and +Dox animals (Fig. 5a). In agreement with this model of PGBD5-induced cell transformation, we observed that endogenous PGBD5 in established G401 and A204 rhabdoid tumor cells was dispensable for cell survival, as assessed using small hairpin RNA (shRNA) interference using two different shRNA vectors, as compared to control shRNA targeting GFP (Fig. 5c & d). Thus, transient expression of *PGBD5* is sufficient to transform cells, as would be predicted from the ability of a catalytically active transposase to induce heritable cellular alterations.

**Fig. 5.**
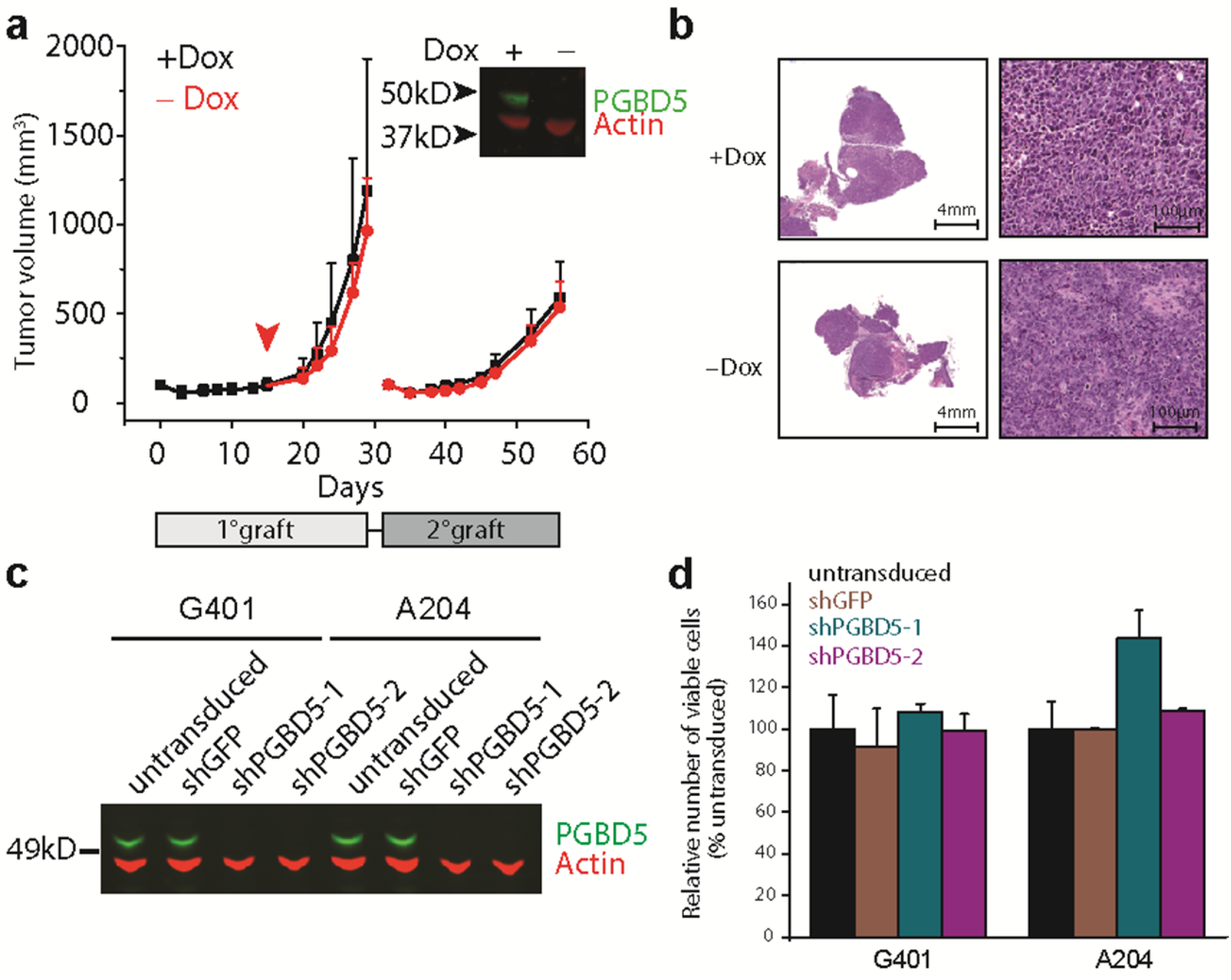
Transient PGBD5 transposase expression is sufficient to transform human cells. Tumor volume of RPE cells as a function of time in primary (light gray box) and secondary (dark gray box) transplants, with *PGBD5* expression induced using doxycycline (black), as indicated. RPE cells were treated with doxycycline *in vitro* for 10 days prior to transplantation. Arrowhead denotes withdrawal of doxycycline from the diet (red). Inset: Western blot of PGBD5 protein, as compared to actin control in cells derived from tumors after primary transplant. **(b)** Representative photomicrographs of hematoxylin and eosin stained tumor sections from doxycycline-inducible PGBD5-expressing RPE tumors after continuous (+Dox) and discontinuous (-Dox) doxycycline treatment. **(c)** Western blot of PGBD5 in G401 and A204 rhabdoid tumor cells upon depletion of *PGBD5* using two independent shRNAs, as compared to non-transduced cells and control cells expressing shGFP. **(d)** Relative number of viable G401 and A204 cells upon 72 hours of *PGBD5* shRNA depletion. Errors bars represent standard deviations of three biological replicates.

### PGBD5-induced transformation requires DNA end-joining repair

If PGBD5-induced cell transformation involves transposase-mediated genomic rearrangements, then this process should depend on the repair of DNA double-strand breaks (DSBs) that are generated by the DNA recombination reactions ^39^. Genomic rearrangements induced by transposases of the DDD/E superfamily involve transesterification reactions that generate DSBs that are predominantly repaired by DNA non-homologous end-joining (NHEJ) in somatic cells ^40^, as is the case for human V(D)J rearrangements induced by the RAG1/2 recombinase ^38^. To test whether PGBD5-induced cell transformation requires NHEJ, we used isogenic RPE cells that are wild-type or deficient for the NHEJ cofactor *PAXX,* which stabilizes the NHEJ repair complex and is required for efficient DNA repair ^41^. In contrast to defects in other NHEJ components, such as *LIG4, PAXX* deficiency does not appreciably alter cell growth or viability but significantly reduces NHEJ efficiency without needing TP53 inactivation to survive ^41^. Thus, we generated RPE cells expressing doxycycline-inducible *PGBD5* that were *PAXX^+/+^* or *PAXX^−/−^,* and confirmed the induction of *PGBD5* and lack of *PAXX* expression by Western immunoblotting (Fig. 6a). Doxycycline-induced expression of PGBD5 in *PAXX^−/−^* but not isogenic *PAXX^+/+^* RPE cells caused the accumulation of DNA damage-associated γH2AX (Fig. 6b, Supplementary Figure 8b), apoptosis-associated cleavage of caspase 3 (Fig. 6c, Supplementary Figure 8a), and cell death (Supplementary Figure 8c). We confirmed the requirement of NHEJ for the repair of PGBD5-induced rearrangements using *Ku80*-deficient mouse embryonal fibroblasts (data not shown). Importantly, PGBD5-mediated induction of DNA damage and cell death in NHEJ-deficient *PAXX^−/−^* cells as compared to the isogenic NHEJ-proficient *PAXX^+/+^* cells was nearly completely rescued by the mutation of D168A/D194A/D386A residues, which are required for transposase activity of PGBD5 (Fig. 6d). Thus, NHEJ DNA repair is required for the survival of cells expressing active PGBD5.

**Fig. 6.**
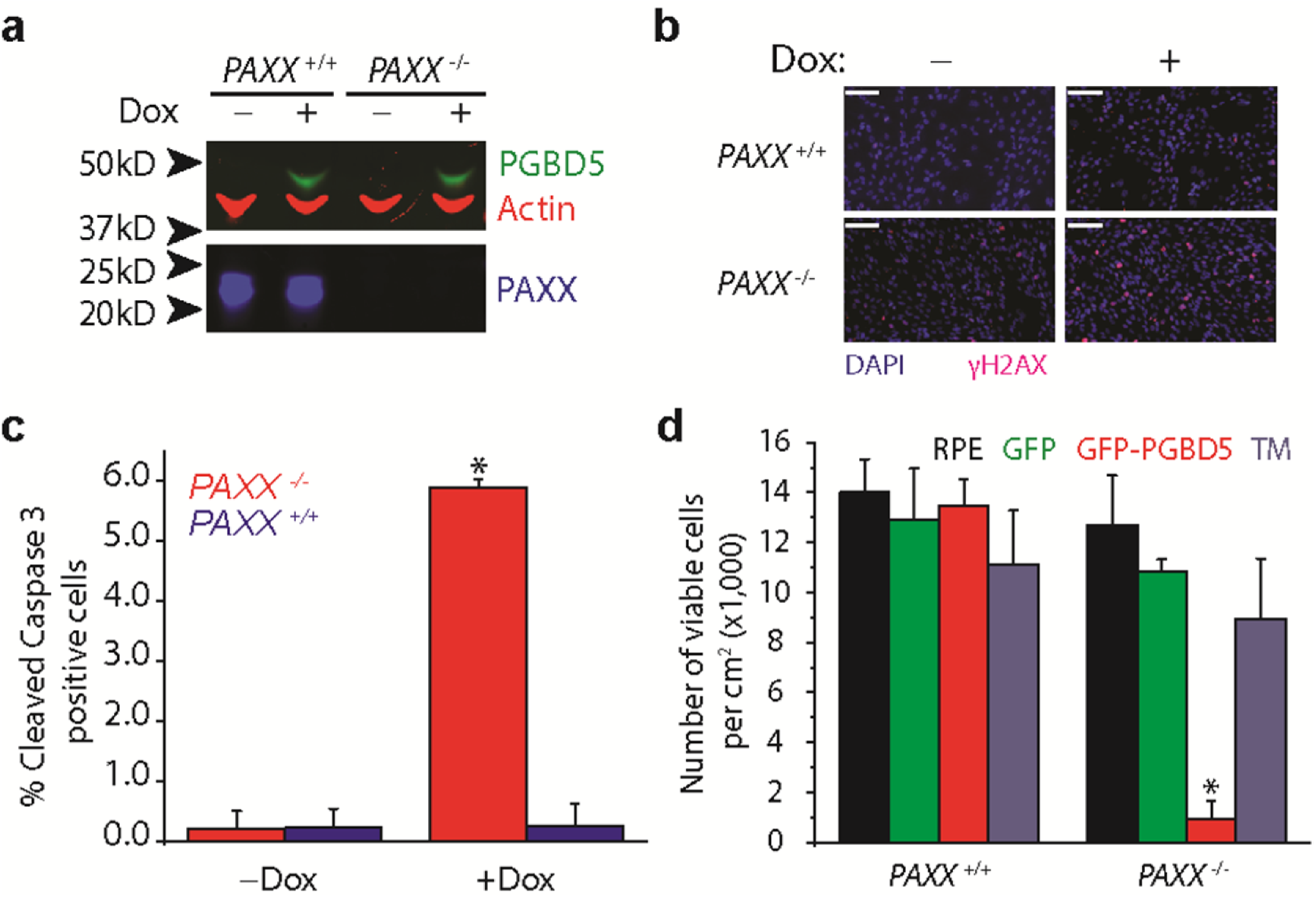
DNA end-joining repair is required for survival of cells expressing active PGBD5. **(a)** Western blot of PGBD5 protein after 24 h of doxycycline (500 ng/ml) treatment of isogenic *PAXX^+/+^* and *PAXX^/-^* RPE cells stably expressing doxycycline-inducible *PGBD5.***(b)** Representative photomicrograph of *PAXX^+/+^* and *PAXX^/-^* RPE cells after 48 h treatment with doxycycline (500 ng/ml) or vehicle control stained for DAPI (blue) and γH2AX(red). Scale bar = 100 μm.**(c)** Fraction of apoptotic cells as measured by cleaved caspase-3 staining and flow cytometric analysis of *PAXX^+/+^* and *PAXX^/-^* RPE cells after treatment with doxycycline or vehicle control. * *p*= 8.7 x 10^-4^ for *PAXX^+/+^* vs. *PAXX^/-^* with doxycycline. **(d)** Number of viable *PAXX^+/+^* and *PAXX^/-^* RPE cells per cm^2^ in monolayer culture as measured by Trypan blue staining after 48 h of expression of *GFP-PGBD5,* as compared to *GFP-PGBD5 D168A/D194A/D386* mutant and GFP-expressing control cells. * *p*= 7.4 x 10^-5^ for *PAXX^/-^GFP-PGBD5* vs. *GFP* control. Error bars represent standard deviations of three biological replicates.

### PGBD5-induced cell transformation involves site-specific genomic rearrangements associated with PGBD5-specific signal sequence breakpoints

The requirements for PGBD5 enzymatic transposase activity, cellular NHEJ DNA repair, and ability of transient *PGBD5* expression to promote cell transformation are all consistent with the generation of heritable genomic rearrangements that mediate PGBD5-induced tumorigenesis. To determine the genetic basis of PGBD5-induced cell transformation, we sequenced whole genomes of PGBD5-induced tumors as well as control GFP-expressing and non-transduced RPE cells, using massively parallel paired-end Illumina sequencing at a coverage in excess of 80-fold for over 90% of the genome (Data S1). As for the rhabdoid tumor genome analysis, we used the assembly-based algorithm laSV as well as conventional techniques (Supplementary Table 3, Supplementary Figs. 9-11, Data S1) ^33,34^. This analysis led to the identification of distinct genomic rearrangements, specifically in PGBD5-induced tumor cell genomes as compared to control GFP and non-transduced RPE cells (Fig. 7a). The identified rearrangements were characterized by intra-chromosomal deletions with a median length of 183 bp, consistent with their apparent limited detectability by conventional genome analysis methods, as well as inversions, duplications and translocations (Supplementary Fig. 12a-c, Data S1). As with genomic rearrangements found in primary human tumors (Fig. 1), the analysis of genomic rearrangements found in PGBD5-transformed RPE cells detected significant enrichment of PSS motifs at the breakpoints of PGBD5-induced tumor structural variants (*p* = 7.2 × 10^−3^, hypergeometric test; Fig. 7b, Data S1). By contrast, breakpoints of structural variants in GFP control RPE cell genomes, presumably at least in part due to normal genetic variation, exhibited no enrichment for PSS motifs (*p* = 0.37). We independently verified these findings using the direct tree graph-based read comparative SMuFin analysis method (Supplementary Fig. 12a, Data S1). In addition, we assessed five of these rearrangements using variant and wild-type allele-specific PCR followed by Sanger DNA sequencing of rearrangement breakpoints to confirm that they are specifically present in PGBD5-transformed but not control GFP-transduced RPE cells (Supplementary Fig. 12d-h). Additionally, we did not find genomic rearrangement breakpoints containing RSS sequences that are targeted by the RAG1/2 recombinase which is not expressed in RPE cells. We also did not find evidence of structural alterations of the annotated human *MER75* and *MER85* piggyBac-like transposable elements, in agreement with the distinct evolutionary history of human *PGBD5* ^30^. Notably, we found that the genomic rearrangements and structural variants observed in PGBD5-induced RPE tumors were significantly enriched for regulatory DNA elements important for normal human embryonal as opposed to adult tissue development (Fig. 7c, Supplementary Table 4).

**Fig. 7.**
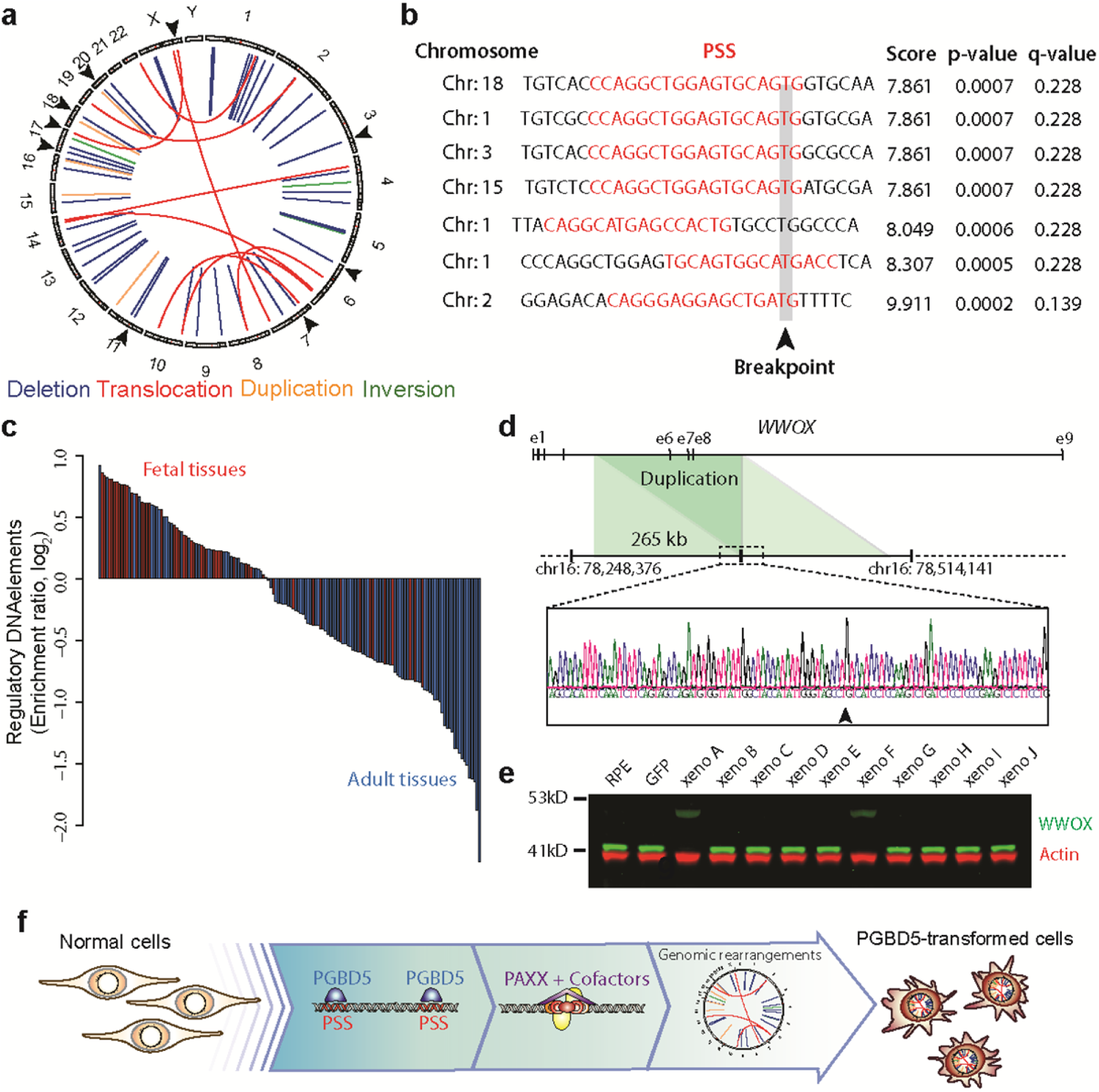
PGBD5-induced cell transformation involves site-specific genomic rearrangements associated with PGBD5-specific signal sequence breakpoints. **(a)** Circos plot of structural variants discovered in RPE-GFP-PGBD5 tumor cells using assembly-based genome analysis. Black arrows on outer circle indicate the presence of PSS at variant breakpoints. **(b)** Representation of 7 breakpoints identified to harbor PSS sequences (red) within 10 bp of the breakpoint junction (arrowhead) of structural variants in PGBD5 expressing RPE cells. Genomic sequence is annotated 5’ to 3’ as presented in the reference genome (+) strand. **(c)** Waterfall plot of enrichment of ENCODE regulatory DNA elements with structural variants in fetal (red) as compared to adult tissues (blue) in *PGBD5*-transformed RPE cells (*p*= 5.7 x 10^-8^).**(d)** Schematic of the *WWOX* gene and its intragenic duplication in GFP-PGBD5-transformed RPE cells (top), with Sanger sequencing chromatogram of the rearrangement breakpoint (bottom). Arrowhead marks the breakpoint. **(e)** Western blot analysis of WWOX in 10 independent GFP-PGBD5-transformed RPE cell tumor xenografts, as compared to control GFP-transduced and non-transduced RPE cells. Actin serves as loading control. **(f)** Schematic model of the proposed mechanism of PGBD5-induced cell transformation, involving association of PGBD5 with genomic PSS sequences, their remodeling dependent on PAXX-meditated end-joining DNA repair, and generation of tumorigenic genomic rearrangements.

To identify genomic rearrangements that may be functionally responsible for PGBD5-induced cell transformation, we analyzed the recurrence of PGBD5-induced genomic rearrangements in 10 different RPE tumors from independent transduction experiments in individual mouse xenografts. We detected 59 PGBD5-induced structural variants per tumor, 42 (71%) of which were deletions, 36 (61%) affected regulatory intergenic elements, with 13 (22%) containing PSS motifs at their breakpoints (Data S1). In particular, we identified recurrent and clonal PSS-associated rearrangements of *WWOX,* including duplication of exons 6-8 (Fig. 7d). *WWOX* is a tumor suppressor gene that controls TP53 signaling ^42^. We confirmed the duplication of exons 6-8 of *WWOX* by PCR and Sanger DNA sequencing (Fig. 7d), and tested its functional consequence on WWOX protein expression by Western immunoblotting (Fig. 7e). Remarkably, this mutation resulted in low level expression of extended mutant form of WWOX protein, associated with loss of wild-type *WWOX* expression, consistent with the dominant negative or gain-of-function activity of mutant *WWOX* in RPE cell transformation. We observed this mutation in 2 out of 10 independent RPE tumors, consistent with its likely pathogenic function in PGBD5-induced cell transformation. To determine its function in PGBD5-induced RPE cells transformation, we depleted endogenous WWOX and ectopically expressed wild-type WWOX in non-transformed wild-type and *WWOX-mutant* PGBD5-induced RPE cell tumors (Supplementary Fig. 13a & d). Consistent with the tumorigenic function of PGBD5-induced mutations of *WWOX*, we found that *WWOX* inactivation was necessary but not sufficient to maintain clonogenicity of PGBD5-transformed RPE tumor cells *in vitro* (Supplementary Fig. 13b-c & e-f). Thus, PGBD5-induced cell transformation involves site-specific genomic rearrangements that are associated with PGBD5-specific signal sequence breakpoints that recurrently target regulatory elements and tumor suppressor genes (Fig. 7f).

## Discussion

We have now found that primary human rhabdoid tumor genomes exhibit signs of PGBD5-mediated DNA recombination, involving recurrent mutations of previously elusive rhabdoid tumor suppressor genes (Fig. 1). These genomic rearrangements involve breakpoints associated with the PGBD5-specific signal (PSS) sequences that are sufficient to mediate DNA rearrangements in rhabdoid tumor cell lines and physical recruitment of endogenous PGBD5 transposase (Fig. 2). The enzymatic activity of PGBD5 is both necessary and sufficient to promote similar genomic rearrangements in primary human cells, causing their malignant transformation (Figs. 3-7).

PGBD5-induced genomic rearrangements comprise a defined architecture, including characteristic deletions, inversions and complex rearrangements that appear distinct from those generated by other known mutational processes. We observe an imprecise relationship of PSS sequences with genomic rearrangement breakpoints, with evidence of incomplete ‘cut-and-paste’ DNA transposition, consistent with potentially aberrant targeting of PGBD5 nuclease activity. While our structure-function studies suggest that PGBD5 induces genomic rearrangements in conjunction with the canonical NHEJ apparatus, it is possible that PGBD5 activity can also promote other DSB repair pathways, such as alternative microhomology-mediated end joining (Supplementary Fig. 14). We confirmed that the catalytic aspartic acid mutants of PGBD5 on average maintain chromatin localization of wild-type PGBD5. It is also possible that these residues contribute to cell transformation due to their interaction with cellular cofactors or assembly of DNA regulatory complexes, or still yet unknown nuclease-independent functions that contribute to cell transformation.

PSS-associated genomic rearrangements induced by PGBD5 in rhabdoid tumors are reminiscent of McClintock’s “mutable loci” induced upon DNA transposase mediated mutations of the *Ds* locus that controls position-effect variegation in maize ^24,43^. Insofar as nuclease substrate accessibility is controlled by chromatin structure and conformation, PGBD5-induced genomic rearrangements indeed may be linked with developmental regulatory programs that control gene expression and specification of cell fate, as suggested by their strong association with developmental regulatory DNA elements. The association of PGBD5-induced rearrangements may involve sequence-specific recognition of human genomic PSS sequences, or alternatively by their accessibility or the presence of cellular co-factors, as determined by cellular developmental states.

Importantly, the spectrum of PGBD5-induced genomic rearrangements and their PSS sequences identified in this study should provide a useful approach to functional characterization of childhood tumor genomes and identification of cancer-causing genomic alterations. In the case of rhabdoid tumors, the association of *SMARCB1* mutations with additional recurrent genomic lesions, such as structural alterations of *CNTNAP2, TENM2* and *TET2* genes that can regulate developmental and epigenetic cell fate specification, may lead to the identification of additional mechanisms of childhood cancer pathogenesis, including those that cooperate with the dysregulation of SWI/SNF/BAF-mediated nucleosome remodeling induced by *SMARCB1* loss. Notably, the recurrence patterns of PGBD5-induced genomic rearrangements in rhabdoid tumors indicate that even for rare cancers, more comprehensive tumor genome analyses will be necessary to define the spectrum of causal genomic lesions and potential therapeutic targets. Similarly, given the existence of distinct molecular subtypes of rhabdoid tumors ^9,10^, it will be important to determine to what extent PGBD5-induced genome remodeling contributes to this phenotypic diversity.

In summary, PGBD5 defines a distinct class of oncogenic mutators that contribute to cell transformation not due to mutational activation but rather as a result of their aberrant induction and chromatin targeting to induce site-specific transforming genomic rearrangements. Our data identify *PGBD5* as an endogenous human DNA transposase that is sufficient to fully transform primary immortalized human cells in the absence of chromosomal instability ^36^. Given the expression of *PGBD5* in various childhood and adult solid tumors, either by virtue of aberrant or co-opted tissue expression, we anticipate that PGBD5 may also contribute to their pathogenesis. Similarly, it will be important to investigate the functions of PGBD5 in normal vertebrate and mammalian development, given its ability to induce site-specific somatic genomic rearrangements in human cells. Finally, the functional requirement for cellular NHEJ DNA repair in PGBD5-induced cell transformation might foster rational therapeutic strategies for rhabdoid and other tumors involving endogenous DNA transposases.

## Methods Summary

A detailed description of the methods is provided as part of the Supplementary Information.

## Supplementary Information

Detailed Materials and Methods, Supplementary Figures 1-14, Supplementary Tables 1-7, and Supplementary Data S1. Genome and chromatin immunoprecipitation sequencing data have been deposited to the NCBI Sequence Read Archive and Gene Expression Omnibus databases (Bioproject 320056 and DataSet GSE81160, respectively). Analyzed data are openly available at the Zenodo digital repository (http://dx.doi.org/10.5281/zenodo.50633).

## Acknowledgments

We are grateful to Alejandro Gutierrez, Marc Mansour, Daniel Bauer, Thomas Look, Hao Zhu, Cedric Feschotte, Michael Kharas, John Petrini and Maria Gil Mir for critical discussions, John Gilbert for editorial support, and Ian MacArthur for technical assistance. This work was supported by the NIH K08 CA160660, P30 CA008748, U54 OD020355, UL1 TR000457, P50 CA140146, Cancer Research UK, Wellcome Trust, Starr Cancer Consortium, Burroughs Wellcome Fund, Sarcoma Foundation of America, Matthew Larson Foundation, Josie Robertson Investigator Program, and Rita Allen Foundation. A.K. is the Damon Runyon-Richard Lumsden Foundation Clinical Investigator.

## Author Contributions

AGH study design and collection and interpretation of the data, RK ChIP-seq, whole genome sequencing and FLEA-PCR data analysis, JZ tumor genome sequencing data analysis with laSV, EJ *in vitro* transformation assays and vector design and cloning, CR *in vitro* transformation assays and vector design and cloning, AE *in vitro* transformation assays and vector design and cloning, ES *in vitro* transformation assays and vector design and cloning, ERF genome sequencing data analysis, SG genome sequencing data analysis, MP genome sequencing data analysis, ANB creation of PAXX deficient cells and study design, CEM genome sequencing data analysis, EDS mouse xenograft study design, MG statistical analysis of datasets, AKE genome sequencing data analysis, MS genome sequencing data analysis, KA genome sequencing data analysis, CRe genome sequencing data analysis, NDS genome sequencing data analysis, EP study design, CRA histological analysis of tumor samples, CWMR study design, HS study design, EM study design, SPJ creation of PAXX-deficient cells and study design, DT genome sequencing data analysis, ZW genome sequencing data analysis, SAA study design, and AK study design, data analysis and interpretation. AK and AGH wrote the manuscript with contributions from all authors.

## Author Information

Correspondence and requests for materials should be addressed to.

## References

1 Vogelstein, B. et al. Cancer genome landscapes. Science 339, 1546–1558, doi:10.1126/science.1235122 (2013).

2 Alexandrov, L. B. et al. Signatures of mutational processes in human cancer. Nature 500, 415–421, doi:10.1038/nature12477 (2013).

3 Cancer Genome Atlas Research, N. et al. The Cancer Genome Atlas Pan-Cancer analysis project. Nature genetics 45, 1113–1120, doi:10.1038/ng.2764 (2013).

4 Futreal, P. A. et al. A census of human cancer genes. Nature reviews. Cancer 4, 177–183, doi:10.1038/nrc1299 (2004).

5 Huether, R. et al. The landscape of somatic mutations in epigenetic regulators across 1,000 paediatric cancer genomes. Nature communications 5, 3630, doi:10.1038/ncomms4630 (2014).

6 Northcott, P. A. et al. Enhancer hijacking activates GFI1 family oncogenes in medulloblastoma. Nature 511, 428–434, doi:10.1038/nature13379 (2014).

7 Mansour, M. R. et al. An oncogenic super-enhancer formed through somatic mutation of a noncoding intergenic element. Science, doi:10.1126/science.1259037 (2014).

8 Molenaar, J. J. et al. Sequencing of neuroblastoma identifies chromothripsis and defects in neuritogenesis genes. Nature 483, 589–593, doi:10.1038/nature10910 (2012).

9 Johann, P. D. et al. Atypical Teratoid/Rhabdoid Tumors Are Comprised of Three Epigenetic Subgroups with Distinct Enhancer Landscapes. Cancer cell 29, 379–393, doi:10.1016/j.ccell.2016.02.001 (2016).

10 Chun, H. J. et al. Genome-Wide Profiles of Extra-cranial Malignant Rhabdoid Tumors Reveal Heterogeneity and Dysregulated Developmental Pathways. Cancer cell 29, 394–406,doi:10.1016/j.ccell.2016.02.009 (2016).

11 Jones, D. T. et al. Dissecting the genomic complexity underlying medulloblastoma. Nature 488, 100–105, doi:10.1038/nature11284 (2012).

12 Fischer, H. P., Thomsen, H., Altmannsberger, M. & Bertram, U. Malignant rhabdoid tumour of the kidney expressing neurofilament proteins. Immunohistochemical findings and histogenetic aspects. Pathology, research and practice 184, 541–547, doi:10.1016/S03440338(89)801499 (1989).

13 Lee, R. S. et al. A remarkably simple genome underlies highly malignant pediatric rhabdoid cancers. The Journal of clinical investigation **In Press**, doi:64400 [pii] 10.1172/JCI64400 (2012).

14 van den Heuvel-Eibrink, M. M. et al. Malignant rhabdoid tumours of the kidney (MRTKs), registered on recent SIOP protocols from 1993 to 2005: a report of the SIOP renal tumour study group. Pediatr Blood Cancer 56, 733–737, doi:10.1002/pbc.22922 (2011).

15 Versteege, I. et al. Truncating mutations of hSNF5/INI1 in aggressive paediatric cancer. Nature 394, 203–206, doi:10.1038/28212 (1998).

16 Roberts, C. W., Leroux, M. M., Fleming, M. D. & Orkin, S. H. Highly penetrant, rapid tumorigenesis through conditional inversion of the tumor suppressor gene Snf5. Cancer cell 2, 415–425, doi:S153561080200185X [pii] (2002).

17 Rousseau-Merck, M. F., Fiette, L., Klochendler-Yeivin, A., Delattre, O. &Aurias, A. Chromosome mechanisms and INI1 inactivation in human and mouse rhabdoid tumors. Cancer genetics and cytogenetics 157, 127–133, doi:10.1016/j.cancergencyto.2004.06.006 (2005).

18 Takita, J. et al. Genome-wide approach to identify second gene targets for malignant rhabdoid tumors using high-density oligonucleotide microarrays. Cancer science 105, 258–264, doi:10.1111/cas.12352 (2014).

19 Smit, A. F. Interspersed repeats and other mementos of transposable elements in mammalian genomes. Curr Opin Genet Dev 9, 657–663 (1999).

20 Kazazian, H. H., Jr. Mobile elements: drivers of genome evolution. Science 303, 1626–1632, doi:10.1126/science.1089670 (2004).

21 Rodic, N. et al. Retrotransposon insertions in the clonal evolution of pancreatic ductal adenocarcinoma. Nat Med 21, 1060–1064, doi:10.1038/nm.3919 (2015).

22 Muotri, A. R. et al. Somatic mosaicism in neuronal precursor cells mediated by L1 retrotransposition. Nature 435, 903–910, doi:10.1038/nature03663 (2005).

23 Shaheen, M., Williamson, E., Nickoloff, J., Lee, S. H. & Hromas, R. Metnase/SETMAR: a domesticated primate transposase that enhances DNA repair, replication, and decatenation. Genetica 138, 559–566, doi:10.1007/s1070901094521 (2010).

24 Hiom, K., Melek, M. & Gellert, M. DNA transposition by the RAG1 and RAG2 proteins: a possible source of oncogenic translocations. Cell 94, 463–470 (1998).

25 Navarro, J. M. et al. Site-and allele-specific polycomb dysregulation in T-cell leukaemia. Nature communications 6, 6094, doi:10.1038/ncomms7094 (2015).

26 Papaemmanuil, E. et al. RAG-mediated recombination is the predominant driver of oncogenic rearrangement in ETV6-RUNX1 acute lymphoblastic leukemia. Nature genetics 46, 116–125, doi:10.1038/ng.2874 (2014).

27 Halper-Stromberg, E. et al. Fine mapping of V(D)J recombinase mediated rearrangements in human lymphoid malignancies. BMC Genomics 14, 565, doi:10.1186/1471216414565 (2013).

28 Dreyer, W. J., Gray, W. R. & Hood, L. The Genetics, Molecular, and Cellular Basis of Antibody Formation: Some Facts and a Unifying Hypothesis. Cold Spring Harb Symp Quant Biol 32, 353–367 (1967).

29 Majumdar, S., Singh, A. & Rio, D. C. The human THAP9 gene encodes an active P-element DNA transposase. Science 339, 446–448, doi:10.1126/science.1231789 (2013).

30 Henssen, A. G. et al. Genomic DNA transposition induced by human PGBD5. eLife 4, doi:10.7554/eLife.10565 (2015).

31 Pavelitz, T., Gray, L. T., Padilla, S. L., Bailey, A. D. & Weiner, A. M. PGBD5: a neural-specific intron-containing piggyBactransposase domesticated over 500 million years ago and conserved from cephalochordates to humans. Mob DNA 4, 23, doi:10.1186/17598753423 (2013).

32 Henssen, A. G. et al. Forward genetic screen of human transposase genomic rearrangements. BMC Genomics 17, 548 (2016).

33 Zhuang, J. & Weng, Z. Local sequence assembly reveals a high-resolution profile of somatic structural variations in 97 cancer genomes. Nucleic acids research 43, 8146–8156, doi:10.1093/nar/gkv831 (2015).

34 Moncunill, V. et al. Comprehensive characterization of complex structural variations in cancer by directly comparing genome sequence reads. Nat Biotechnol 32, 1106–1112, doi:10.1038/nbt.3027 (2014).

35 Bralten, L. B. et al. The CASPR2 cell adhesion molecule functions as a tumor suppressor gene in glioma. Oncogene 29, 6138–6148, doi:10.1038/onc.2010.342 (2010).

36 Hahn, W. C. et al. Creation of human tumour cells with defined genetic elements. Nature 400, 464–468,doi:10.1038/22780 (1999).

37 Landree, M. A., Wibbenmeyer, J. A. & Roth, D. B. Mutational analysis of RAG1 and RAG2 identifies three catalytic amino acids in RAG1 critical for both cleavage steps of V(D)J recombination. Genes & development 13, 3059–3069 (1999).

38 Lu, C. P., Posey, J. E. & Roth, D. B. Understanding how the V(D)J recombinase catalyzes transesterification: distinctions between DNA cleavage and transposition. Nucleic acids research 36, 2864–2873, doi:10.1093/nar/gkn128 (2008).

39 Gellert, M. V(D)J recombination: RAG proteins, repair factors, and regulation. Annu Rev Biochem 71, 101–132, doi:10.1146/annurev.biochem.71.090501.150203 (2002).

40 Mitra, R., Fain-Thornton, J. & Craig, N. L. piggyBac can bypass DNA synthesis during cut and paste transposition. The EMBO journal 27, 1097–1109, doi:10.1038/emboj.2008.41 (2008).

41 Ochi, T. et al. DNA repair. PAXX, a paralog of XRCC4 and XLF, interacts with Ku to promote DNA double-strand break repair. Science 347, 185–188, doi:10.1126/science.1261971 (2015).

42 Aldaz, C. M., Ferguson, B. W. & Abba, M. C. WWOX at the crossroads of cancer, metabolic syndrome related traits and CNS pathologies. BiochimBiophysActa 1846, 188–200, doi:10.1016/j.bbcan.2014.06.001 (2014).

43 McClintock, C. B. The origin and behavior of mutable loci in maize. Proceedings of the National Academy of Sciences of the United States of America 36, 344–355 (1950).

